# Proteomic and metabolomic profiling underlines the stage- and time-dependent effects of high temperature on grape berry metabolism

**DOI:** 10.1101/810481

**Authors:** David Lecourieux, Christian Kappel, Stéphane Claverol, Philippe Pieri, Regina Feil, John E. Lunn, Marc Bonneu, Lijun Wang, Eric Gomès, Serge Delrot, Fatma Lecourieux

## Abstract

Climate change scenarios predict an increase in mean air temperatures and in the frequency, intensity, and length of extreme temperature events in many wine-growing regions worldwide. Because elevated temperature has detrimental effects on the berry growth and composition, it threatens the economic and environmental sustainability of wine production. Using Cabernet Sauvignon fruit-bearing cuttings, we investigated the effects of high temperature (HT) on grapevine berries through a label-free shotgun proteomic analysis coupled to a complementary metabolomic study. Among the 2279 proteins identified, 592 differentially abundant proteins were found in berries exposed to HT. The gene ontology categories “Stress”, “Protein”, “Secondary metabolism” and “Cell wall” were predominantly altered under HT. High temperatures strongly impaired carbohydrate and energy metabolism, and the effects depended on the stage of development and duration of treatment. Transcript amounts correlated poorly with protein expression levels in HT berries, highlighting the value of proteomic studies in the context of heat stress. Furthermore, this work reveals that HT alters key proteins driving berry development and ripening. Finally, we provide a list of differentially abundant proteins that can be considered as potential markers for developing or selecting grape varieties that are better adapted to warmer climates or extreme heat waves.

## INTRODUCTION

Grapevine is economically the most important fruit crop in the world, providing dried fruits, table grapes and the basis for wine making (www.oiv.int; www.fao.org). The combination of climate, soil and viticultural practices in a given region constitutes a highly interactive system known as “terroir” (Deloire et al. 2005). The interaction between “terroir” and the plant’s genotype (cultivar) significantly affects vine development and berry composition, and thereafter determines wine quality and character (Van Leeuwen and Seguin 2006). Grapevine requires suitable temperatures, radiation intensities and duration, and water availability during the growth cycle for optimal fruit production and quality (Kuhn et al. 2014). Because of on-going climate change, the environmental parameters in most wine producing regions around the world are expected to change, potentially lowering productivity and altering the traditional characteristics (“typicity”) of the wine from a given region (Schultz and Jones 2010). While some of these climatic constraints can be overcome, at least partially, by viticultural practices, temperature remains more difficult to control under field conditions (Van Leeuwen et al. 2013). Climate change scenarios predict an increase in global mean temperatures between 1.5 and 5°C, during the 21^th^ century (IPCC, 2014), in addition to an increase in the frequency, intensity, and duration of extreme temperature events (Lobell et al. 2008). Considering that air temperature is a key climate variable affecting grapevine phenology, berry metabolism and composition (Jones et al. 2005; Martínez-Lüscher et al. 2016), global warming represents a serious threat for wine production and/or typicity in many wine-growing regions worldwide (Fraga et al. 2016).

Temperatures above the optimum are sensed as heat stress by living organisms. Heat stress disturbs cellular homeostasis and can lead to severe retardation in growth and development and even death (Wahid et al. 2007). In grapevine, warming promotes vegetative growth and disturbs the plant’s carbon balance (Greer and Weedon 2012), adversely affecting flower set and young berry development (Greer and Weston 2010). HS also impacts fruit primary and secondary metabolism, desynchronizing sugar and organic acid metabolism and delaying sugar and polyphenol accumulation during ripening (Torregrosa et al. 2017; Gouot et al. 2018). As a consequence, heat waves occurring during key berry development stages can dramatically affect the organoleptic properties of wine even when more favourable weather conditions prevail for the remaining of the season.

In this context, the common viticultural practice of removing leaves around the fruit zone to enhance aroma development and avoid fungal infections (Alem et al. 2018), might be questioned since it affects light exposure and temperature of the berries. Because of the direct irradiative effects, temperature within the fruit may differ very significantly from that of the whole plant or surrounding air (Cola et al., 2009). The microclimate (mostly temperature and light conditions) strongly influences the metabolite composition of berries, affecting flavonols, anthocyanins, amino acids and aroma precursors (Downey et al. 2006; Pereira et al. 2006; Asproudi et al. 2016; Young et al. 2016; Reshef et al. 2017). These effects can be explained, at least in part, by the substantial remodelling of the fruit transcriptome observed under these conditions (du Plessis et al. 2017; Lecourieux et al. 2017).

In a previous report, we showed dramatic biochemical and transcriptomic changes in heat-exposed berries, depending on both the developmental stage and the stress duration (Lecourieux et al. 2017). Although transcriptomic analyses have offered comprehensive insights into the mechanisms underlying the HS responses in grape (Liu et al. 2012; Carbonell-Bejerano et al. 2013; Rienth et al. 2014b; Jiang et al. 2017; Lecourieux et al. 2017), it is recognised that changes in transcript abundance do not always correlate with the levels of the encoded proteins (Keller and Simm 2018). Furthermore, proteins represent a more functional level of gene expression than transcripts because they have a direct impact on metabolism and other cellular processes. Therefore, state of the art mass spectrometry analysis of the proteome is a powerful tool to study molecular mechanisms and biological traits in plants, including responses to abiotic stresses (Kosova et al. 2018). Using shotgun proteomic tools such as iTRAQ (isobaric Tags for Relative and Absolute Quantitation) and label-free quantification techniques, several proteomic studies have recently investigated the heat responses of grapevine leaves (Liu et al. 2014; Jiang et al. 2017) or grape cell cultures (George et al. 2015).

To better understand the consequences of high temperature (HT), inherent to leaf removal, on berry development and to improve our knowledge about the molecular mechanisms involved in grapevine fruit response to HT, the present work combines transcriptomic, proteomic and metabolite analysis using Cabernet Sauvignon fruit-bearing cuttings. A label-free proteomic approach was used to investigate changes in the grape berry proteome resulting from direct exposure to elevated temperature. The collected proteomic data were compared with our previous transcriptomic analysis performed on the same berry samples (Lecourieux et al., 2017). These data showed a strong and negative effect of HT on proteins related to carbohydrate and energy metabolism, which led us to analyse the impact of HT on primary metabolites using tandem mass spectrometry (LC-MS/MS). Finally, we focused on proteins accumulating in berries exposed to HT that are potentially involved in thermotolerance. Altogether, this work helps to better understand the consequences of heat stress on developing grape berries and provides a list of potential heat-tolerance molecular markers that could be useful for breeders and viticulturists.

## RESULTS AND DISCUSSION

### Quantitative analysis of heat stress responsive proteins in grape berries

Experiments were performed on fruit-bearing cuttings of *V. vinifera* Cabernet Sauvignon as described in Lecourieux et al. (2017). In brief, berries at three different stages of development were exposed to elevated temperatures for 12 h (7:00 am to 7:00 pm) during each day of treatment and harvested for analysis after 1 and 7 days of treatment. The average pulp temperature of the heat-treated berries was about 8°C higher than in control berries. A label-free proteomic approach was performed to identify proteins with differential abundance between control and heat-treated berry samples.

Using the parameters and filters described in the Material and Methods, the conducted label-free experiment led to the identification of 3972 protein groups (from here on referred to as proteins) in the combined dataset from the six conditions (three stages x two treatment durations), of which 2279 showed two or more unique peptides (**Table 1**). Protein abundance changes between control and heat-exposed fruits were considered to be significant when a minimum fold change of 1.5 (log_2_FC > 0.58 or < −0.58) was reached, with a significance threshold of *p* < 0.05. Only proteins identified in all three biological replicates were considered for quantification. Among the quantified proteins with a minimum of two unique peptides, 592 DAPs (differentially abundant proteins) displayed significant differences in abundance between control and HT berries in at least one of the six conditions. The entire dataset is available via ProteomeXchange (identifier PXD014693) and in Supplementary **Table S1.** Further information on the 592 DAPs is provided in Supplementary **Table S2**. Among these 592 DAPs, 501 proteins were identified in only one condition (139 up-regulated and 362 down-regulated) while the 91 remaining proteins were retrieved in more than one condition (16 up-regulated, 61 down-regulated, and 14 proteins that were up-regulated in some conditions but down-regulated in others) (**Table 1 and Table S2**). More DAPs were identified at the ripening stage (304 DAPs) than at the green (146 DAPs) or veraison stages (222 DAPs). Whereas the numbers of up- and down-regulated proteins were similar at veraison (115 up, 107 down), more down-regulated proteins were found when the HT was applied at the green stage (24 up, 122 down) or ripening (49 up, 255 down) stages (**Figure 1A**). The DAPs identified after short (1 day) treatments were mostly different from those observed after long (7 days) treatments. Indeed, only 1 (up), 9 (4 up, 5 down) and 18 (2 up, 16 down) common DAPs between 1D or 7D of HT were observed at the green, veraison and middle ripening stages, respectively (**Figure 1B and Tables S2, S3, S4 & S5**). Similarly, at all three stages, only one common down-regulated DAP was identified after 1 day of HT (VIT_12s0057g01020, fasciclin-like protein), with none being detected after 7 days of treatment (**Table S2**). The number of common DAPs per stage increased when looking for proteins with modified abundance after 1 and/or 7 days treatment. In this case, 7 DAPs (3 up, 4 down) were found in common for the three stages, and up to 31 DAPs were obtained when comparing heat effects between green and veraison stages (**Figure 1C and Supplementary Table 2**).

**Figure 1.**
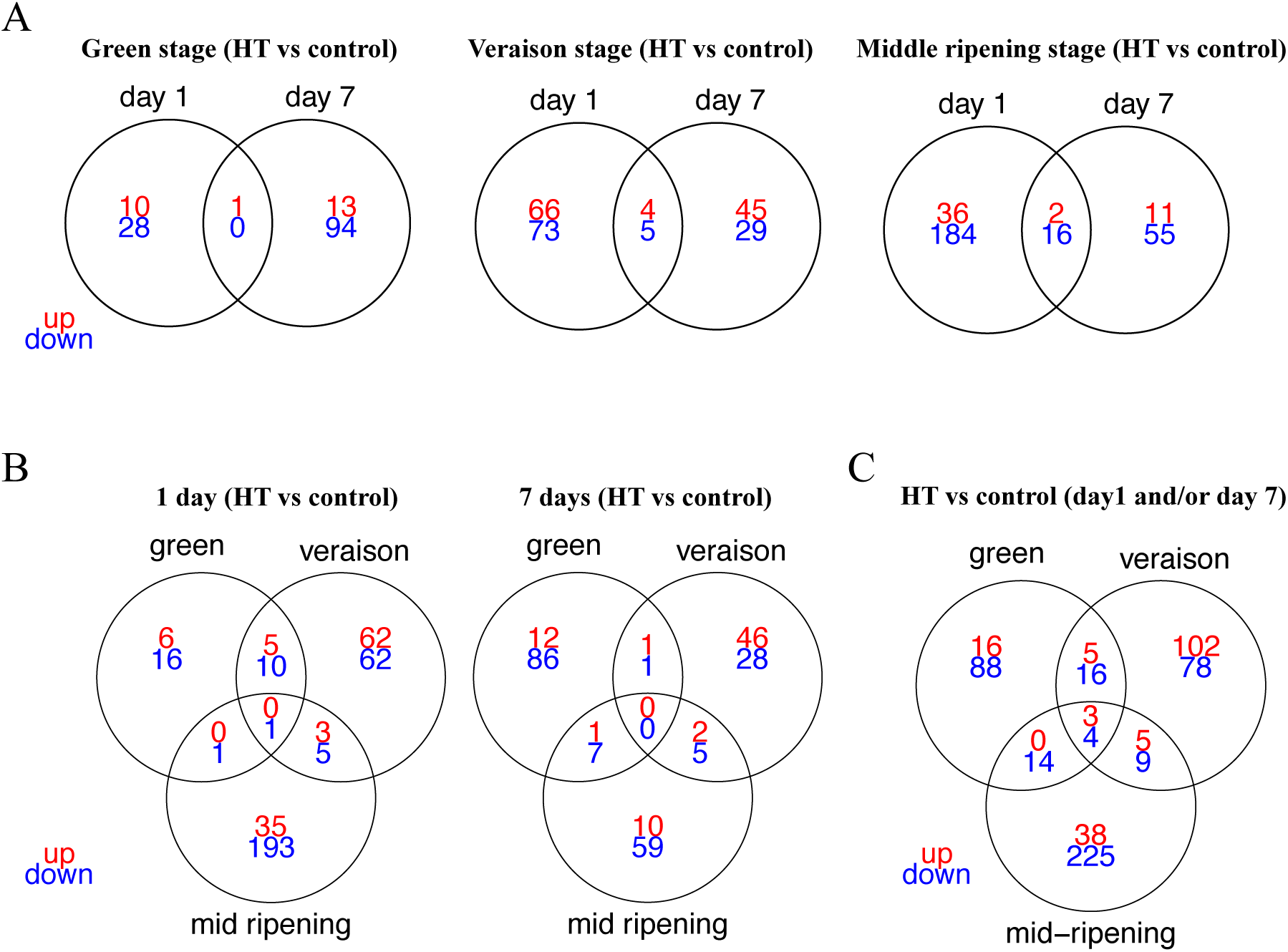
Venn diagrams displaying the numbers of DAPs in heat-treated berries, according to the developmental stage (A) or the treatment duration (B). DAPs were selected using a 1.5 fold expression change and an adjusted p < 0.05 (with FDR correction). Only proteins identified in all three biological replicates and with a minimum of two unique peptides were considered for quantification. The detailed information of the common DAPs upon HT for each stage or for each exposure time was listed as in Supplementary Tables 1 and 2.

**Table 1:**
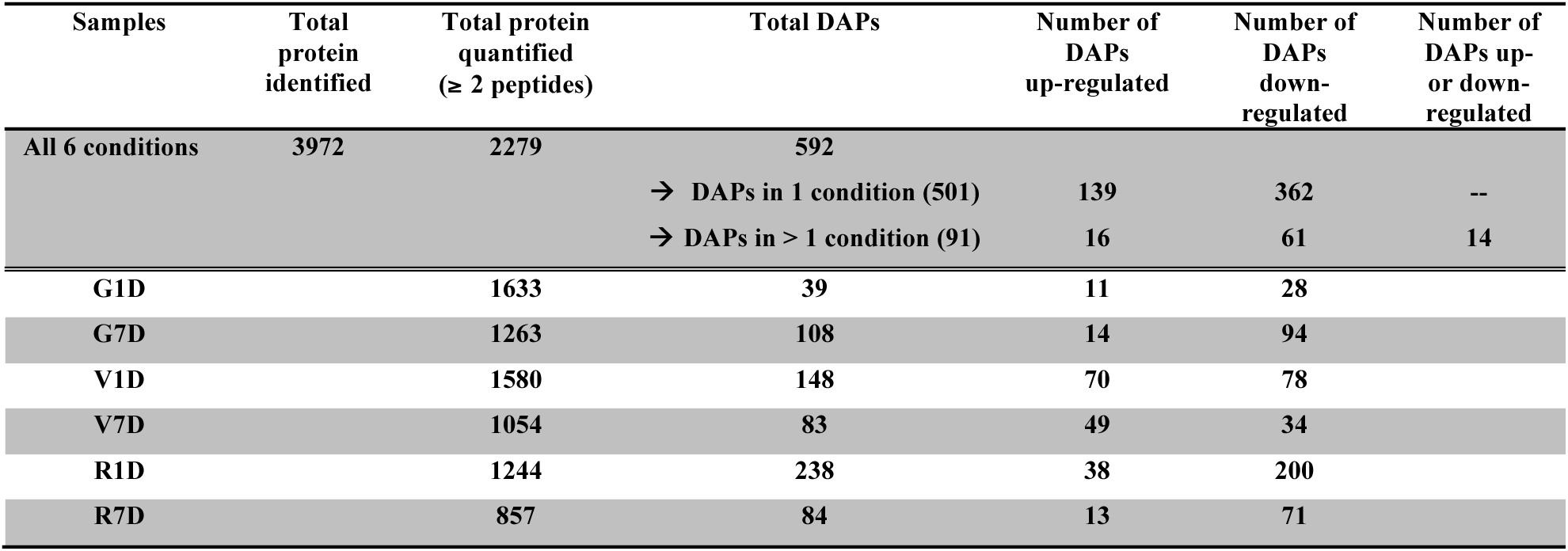
Amount of identified proteins and differentially abundant proteins in control and heat berries according to the developmental stage and the stress duration

To determine the functional categories affected by HT, the identified DAPs were functionally classified using MapMan BIN codes (Usadel et al. 2005) and a *V. vinifera* mapping file (Lecourieux et al., 2017). The predominant proteins changed in berries in response to HT were assigned to “protein metabolism” (114/592 DAPs, 19.3%), “stress” (54/592 DAPs, 9.1%), “secondary metabolism” (31/592 DAPs, 5.2%) and “cell wall” (25/592 DAPs, 4.2%) (**Figure 2**). Except for the categories “stress” (36 up, 18 down) and “RNA” (10 up, 9 down), all the others categories contained more down-regulated proteins that up-regulated ones, with particularly asymmetrical changes in some categories, such as “lipid metabolism” (0 up, 16 down) and “transport” (2 up, 17 down). No particular effect of HT according to the developmental stage was observed on these functional categories, whereas categories linked to carbohydrate and energy metabolism (glycolysis, tricarboxylic acid (TCA) cycle, fermentation, gluconeogenesis, minor and major carbohydrate (CHO) metabolism) or related to “photosynthesis”, “cell wall” and “redox” were mostly impacted during veraison and/or ripening stages (**Supplementary Table 2**).

**Figure 2.**
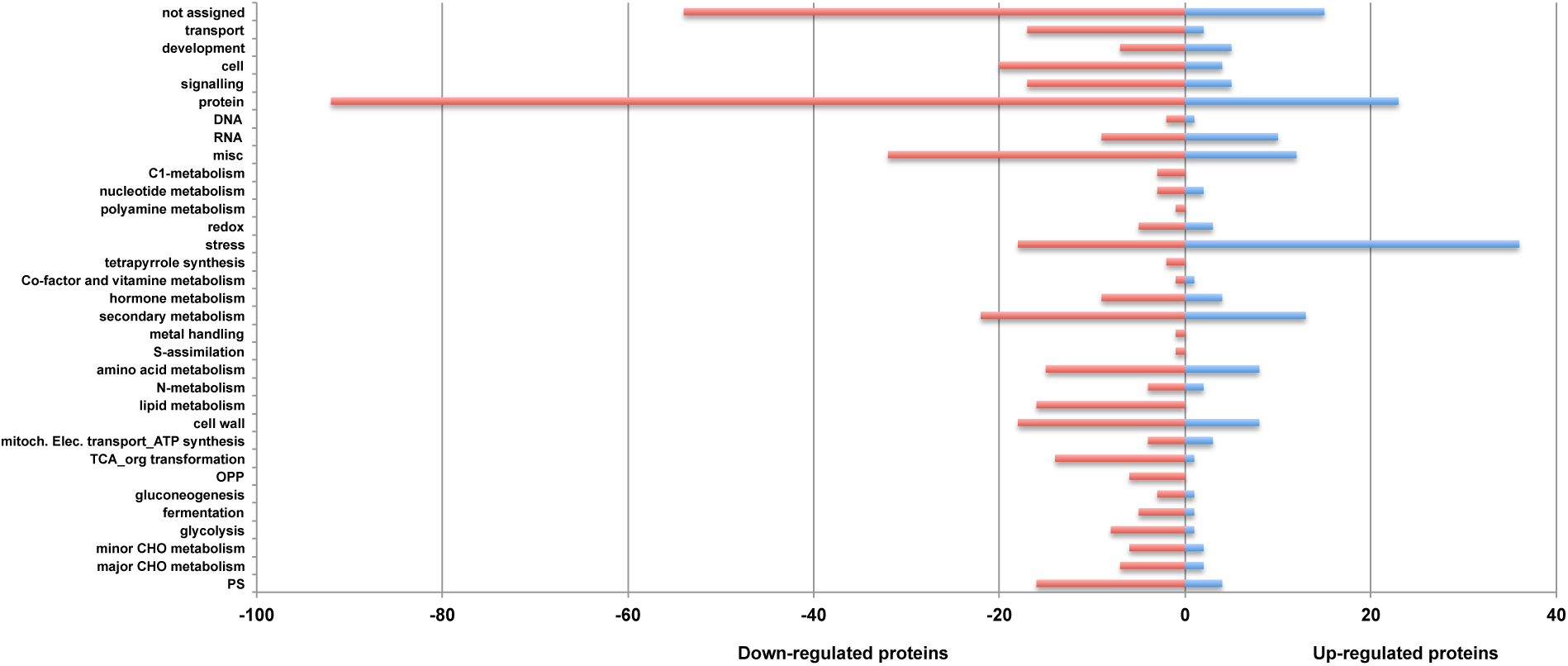
Total number of differentially abundant proteins for each MapMan main category. Category description is given on the y-axis. Numbers of down- and up-regulated proteins across the 6 experimental conditions are provided on the x-axis for each category.

### Poor correlation between the transcriptome and proteome in heat-stressed berries

Global analyses based on gene expression and protein profiles provide a powerful tool for understanding how organisms respond to environment changes. To integrate proteomic data with transcriptomic results, the present protein abundance data were compared to each corresponding differentially expressed gene (DEG) identified in the same grape berry samples exposed to HT (Lecourieux et al., 2017). Whereas 592 DAPs were retrieved in at least one of the six HT conditions (**Table 1, Supplementary Table S1**), 6848 DEGs were identified using the same berry samples (Lecourieux et al. 2017). A linear correlation between differentially expressed transcripts and the corresponding proteins was established for 64 out of 592 DAPs **(Figure 4, Supplementary Table S11)**. Two types of correlation profiles were found. The first one corresponds to proteins and transcripts that exhibit profiles with the same tendency (up- or down-regulation in response to HT) and the second one includes proteins and transcripts that are inversely correlated for a same gene. The best relationship between mRNA and protein levels (39 DAPs) was observed when HT was applied for one day at veraison, and for proteins belonging to functional BINs associated to “heat stress” (25 members), “hormone” (4), and “secondary metabolism” (5) **(Figure 4, Supplementary Table S11)**.

**Figure 3.**
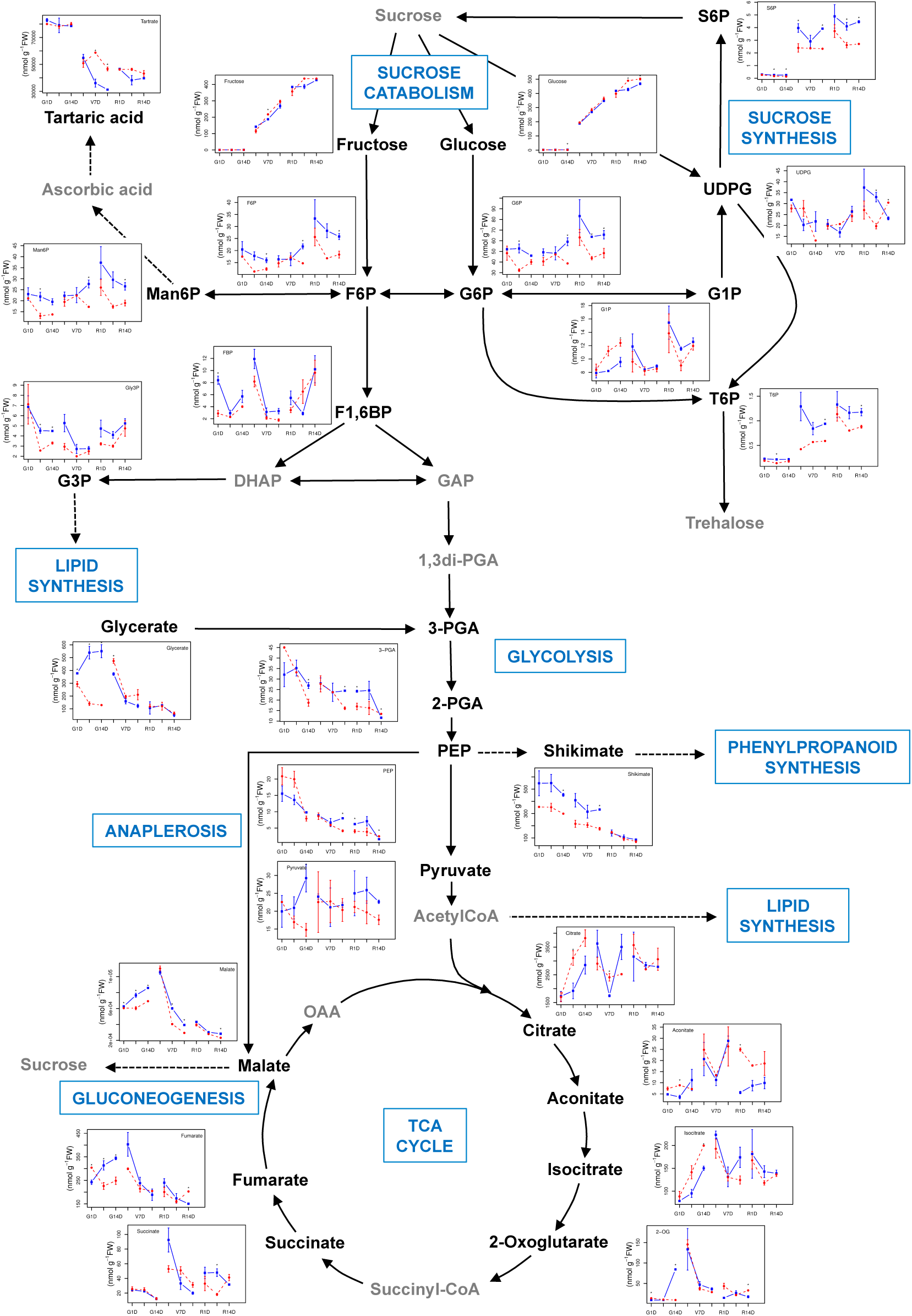
Metabolite profiles of Cabernet Sauvignon developing berries exposed (red line) or not (blue line) to heat stress. Metabolites are shown in their respective metabolic pathways (sugar metabolism, glycolysis, and the TCA cycle) and their temporal profiles during berry treatment (µmol.g^−1^ FW for glucose, and fructose; and nmol.g^−1^ FW for the others) are presented alongside. For each profile, the x-axis shows the developing stage (G: middle-green, V: véraison and R: middle-ripening) and the stress duration (1, 7 and 14 days).

**Figure 4.**
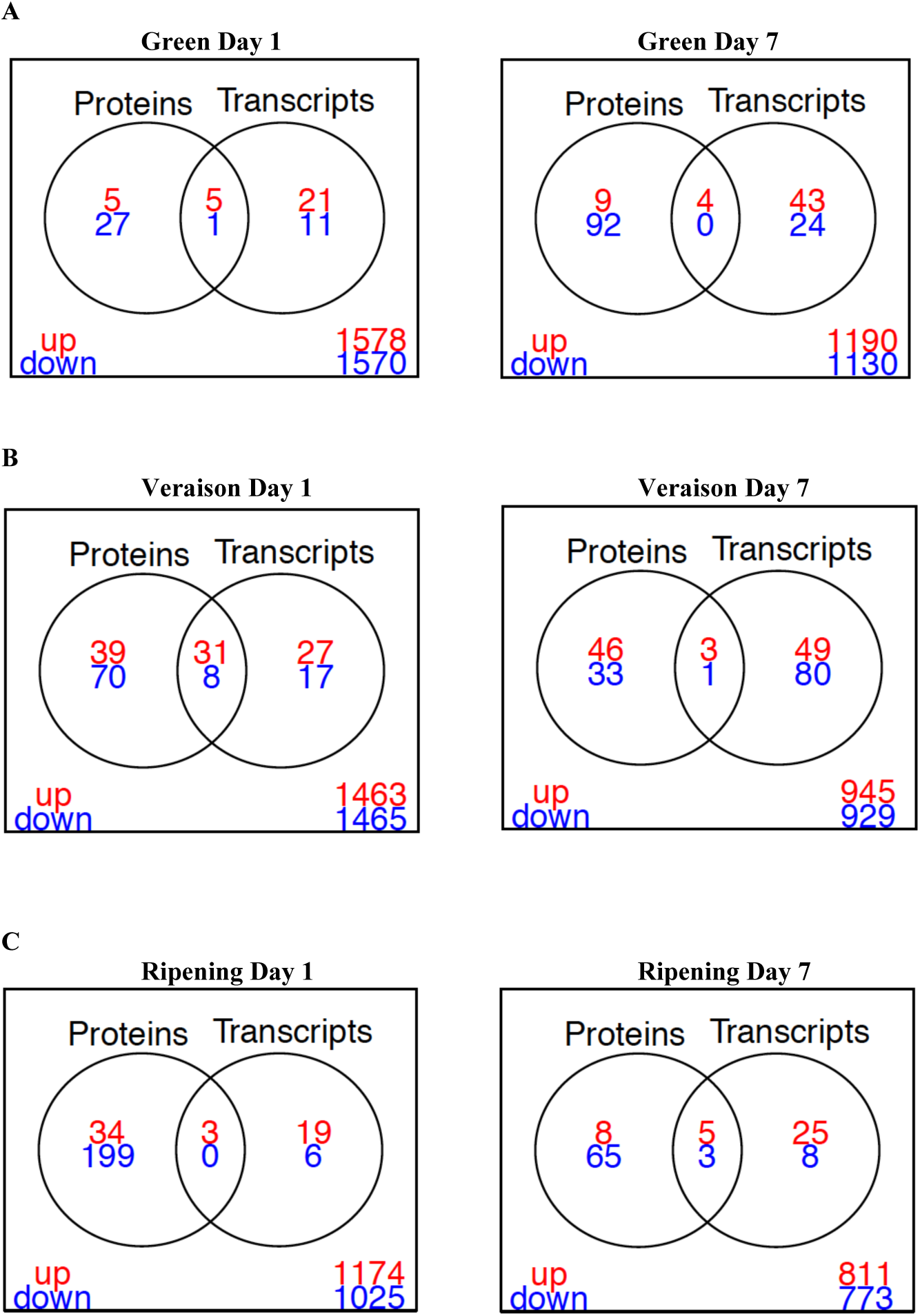
Correlation between the expression of proteins and mRNAs in developing control and HT berries. DAPs and DEGs were selected using a 1.5 and 2 fold expression change, respectively, and an adjusted p < 0.05 (with FDR correction). Venn diagrams display the number of DAPs and DEGs (red: up-regulated and blue: down-regulated) according to the developing stage (A: middle-green, B; véraison and C: middle-ripening) and the stress duration (1 and 7 days).

Our results underline the poor correlation between the proteome and the transcriptome in HT berries, with only 11% (64/592 DAPs) of the differentially abundant proteins that were also impacted at the mRNA expression level. This is not unexpected since HT leads to denaturation and aggregation of proteins, and because translation, especially under elevated temperature, is governed by a variety of regulatory mechanisms independent of transcription rate (Bokszczanin and Fragkostefanakis 2013). These include the impacts of miRNAs on mRNA stability and translatability, mRNA half-life, global and transcript specific translational rates, as well as protein turnover. Any lag between the synthesis of a gene transcript and its translation into a protein could also give rise to an apparent discrepancy between transcript and protein abundance at a given sampling time (Keller and Simm 2018). Last but not least, the poor correlation between transcript and protein levels in heat-treated berry samples may also be due to alternative splicing (AS) events. Increasing evidence has shown that AS is a critical post-transcriptional event and plays an important role in plant stress responses (Baurle 2016). Particularly, many genes are under transcriptional regulation by the circadian clock, and several circadian clock genes show AS in response to even small changes in temperature (James et al. 2018). In grapevine, AS changes were observed for nearly 70% of the genes in leaves exposed to temperatures above 35°C. In the same study, only 64 out of the 808 identified DAPs were retrieved as differentially expressed at the transcript level (Jiang et al. 2017), that fits well with the weak transcript/protein correlation observed in our work. Jiang et al. (2017) suggested that AS changes under high temperature could provide molecular plasticity allowing the plants to adapt to stress conditions. Using Arabidopsis (*Arabidopsis thaliana*) plants exposed or not to a non-lethal HT, referred to as priming, Ling et al. (2018) have recently demonstrated that AS functions as a novel component of heat-shock memory.

Despite the generally poor correlation between individual transcripts and proteins levels in berries exposed to HT, there was a much better correspondence at the functional category level, with transcripts and proteins assigned to the “stress”, “protein metabolism” and “secondary metabolism” MapMan BINs showing consistent responses as a group (**Supplementary Figure S1**). We also noticed that, at the transcript level, most of the functional categories were differentially affected according to the developmental stage and the stress duration whereas it was less obvious at the protein level. The best relationship between transcripts and proteins levels was observed with genes related to the category “stress”, and especially to the “heat” sub-category. Among the 30 HSPs whose protein abundance increased in response to HT, 25 were also up-regulated at the transcript level (**Supplementary Table S12**). Most proteins are inherently thermo-labile, with some becoming denatured at the temperatures that grapevines experience during a heat wave. In addition, high temperature is often associated with oxidative stress leading to production of reactive oxygen species that irreversibly damage proteins. Maintaining proteins in stable and functional conformations and preventing their aggregation are essential for survival of cells under high temperature. HSPs function as molecular chaperones in maintaining protein quality and folding, and are required for the acquisition of plant thermotolerance (Bokszczanin and Fragkostefanakis 2013). It is well established that transcription factors from the HSF family regulate the heat stress response in plants, through a massive up-regulation of *HSP* genes (Scharf et al. 2012). However, as transcription factors are low abundance proteins they largely escaped proteomic detection, and no HSF was retrieved in the present study despite the observed transcriptional up-regulation of six *VviHSF* genes in heat-exposed berries (Lecourieux et al. 2017).

In good agreement with our transcriptomic data, the protein abundance of the functional category “Protein metabolism” was strongly impacted with nearly 20% of the identified DAPs that belong to this category being affected, especially those belonging to the sub-categories “protein synthesis” (34 DAPs) and “protein degradation” (55 DAPs) (**Supplementary Table S6**). However, the HT triggered a strong down-accumulation effect on the “protein metabolism” category (92 out of the 114 DAPs), in contrast with what was observed on transcripts (333 up- and 307 down-regulated genes). A number of studies demonstrated that different abiotic stresses, including heat, result in a general inhibition of protein translation (Yanguez et al. 2013). Accordingly, we observed a significant decrease in abundance of many proteins related to protein synthesis including various ribosome structural proteins and several translation initiation factors (**Supplementary Table S6**). Whereas inhibiting translation under stress helps to reduce energy consumption, certain specific mRNAs are selectively translated to produce relevant proteins involved in the proper establishment of the stress adaptation process. By studying the translatome of Arabidopsis seedling exposed to HT, Yanguez et al. (2013) observed that regulation of gene expression at the translational level is superimposed on regulation at the transcriptional level, and often reinforces the response of individual genes/proteins to heat stress. Protein translation is accomplished by the combined actions of the ribosomes and ancillary proteins such as initiation, elongation, and termination factors. Yanguez et al. (2013) showed that translation could be regulated at the initiation phase in Arabidopsis seedlings exposed to HT, even if the mechanisms involved in this regulation remain unclear. To date, there is some evidence for the role of different plant translation initiation factors (eIFs) in various abiotic stresses responses (Dutt et al. 2015). Our proteomic data highlighted the increase in abundance of four eIFs upon HS, namely eIF1A (VIT_15s0046g01500), two isoforms of eIF3 (VIT_14s0060g01630, VIT_00s0880g00020) and eIF5 (VIT_14s0006g01990), whereas others eIF isoforms were decreased (**Supplementary Table S6**). eIF5 (VIT_14s0006g01990) showed the highest accumulation score (Log_2_FC 5.22) within this functional category. The importance of eIF5 proteins in thermotolerance was highlighted in pea (Suragani et al. 2011) and Arabidopsis (Xu et al. 2011; Zhang et al. 2017).

### Heat stress alters key proteins driving berry development and ripening

#### Heat stress impacts the secondary metabolism in developing berry

The strong effects of HT observed on transcripts assigned to the “secondary metabolism” functional category (291 DEGs across the six conditions, (Lecourieux et al. 2017)) were confirmed at the protein level (31 out of the 592 DAPs), albeit to a lesser extent (**Supplementary Table S8**). These 31 DAPs (11 up, 20 down) mainly belonged to the subcategories: “isoprenoids” (7 DAPs), “phenylpropanoid-lignin” (12) and “flavonoids” (7).

HT altered expression of numerous genes related to aromatic potential, including genes encoding biosynthetic enzymes for terpenes and methoxypyrazines (Lecourieux et al. 2017), but the proteomic data from the current study suggested that these changes in transcript abundance were not translated through to the protein level. A few proteins related to isoprenoid subfamily were down-accumulated under HT including a 3-ketoacyl-CoA synthase (KCS10, VIT_04s0008g02250), which is involved in the synthesis of very long chain fatty acids that are incorporated into plant lipids (precursors of cuticular waxes), and several enzymes related to the synthesis of precursors of terpenoids (**Supplementary Table S8**). By contrast, a carotenoid cleavage dioxygenase (CCD1, VIT_13s0064g00810) was found more abundant in stressed berries at veraison. Expression of various *VvCCD* genes was reported to increase towards berry ripening and enzymes in this family were demonstrated to promote desirable flavor and aroma compounds in grapes and wine (Mathieu et al. 2005; Rienth et al. 2014a; Alem et al. 2018). The putative role of CCD proteins in heat responses is still unknown but these proteins were suggested to enhance tolerance to various abiotic stresses through their enzymatic products (e.g. apocarotenoids) acting as stress signals (Havaux 2014). For instance, some members of the CCD family are involved in generating carotenoid-derived phytohormones, such as strigolactones known to be involved in various plant developmental and adaptation processes (Jia et al. 2018).

It is well known that extreme high temperatures significantly affect the phenylpropanoid content of grape berries, with the impact being dependent on the intensity and duration of the HS as well as the developmental stage of the fruit (Gouot et al. 2018). Consistent with changes observed at the transcript level (Lecourieux et al. 2017), HT triggered an enhanced abundance of proteins involved in lignin synthesis, including shikimate *O*-hydroxycinnamoyltransferase (HCT; VIT_09s0018g01190), cinnamyl alcohol dehydrogenase (CAD; VIT_03s0180g00260, VIT_02s0025g03100, VIT_00s0615g00030) and caffeic acid O-methyltransferase (CCoAOMT1; VIT_16s0098g00850). A significant increase in lignin biosynthesis would confer additional strength to lignified secondary cell wall, helping berries to protect against shriveling mediated-HT. For instance, the activation of the lignin pathway is a reaction to advanced stages of dehydration in grape berry fruit (Zamboni et al. 2010). Key enzymes of the general phenylpropanoid pathway (phenylalanine ammonia-lyase PAL, VIT_06s0004g02620, VIT_08s0040g01710, VIT_13s0019g04460; cinnamate 4-hydroxylase C4H, VIT_06s0004g08150; 4-coumarate-CoA ligase 4CL; VIT_11s0052g01090), enzymes from the flavonoid biosynthesis pathway (dihydroflavonol 4-reductase DFR, VIT_18s0001g12800; flavonoid 3’-hydroxylase F3’H, VIT_17s0000g07210; flavonoid 3’,5’-hydroxylase F3’5’H, VIT_06s0009g02860; flavanone 3’-hydroxylase F3H, VIT_18s0001g14310; isoflavone reductase, VIT_03s0038g04700) as well as enzymes mediating the glycosylation of flavonoids (UDP-glucosyl transferases, VIT_12s0055g00290, VIT_02s0033g00130, VIT_17s0000g04750; UDP-rhamnosyl transferase, VIT_00s0218g00150) were also affected at the protein level upon HT. Most of these proteins were less abundant in heated berries, suggesting a sharp decline of the corresponding enzymatic activities (**Supplementary Tables S2 & S8**). This lower amount of key enzymes probably contributes to the reduction of flavonoid synthesis in grape berries exposed to heat. However, other concomitant processes may explain the decrease of flavonoid contents in warmed berries, including reduced biosynthetic enzyme activities due to non-optimal temperature, enzymatic oxidation or chemical degradation after ROS scavenging (Gouot et al. 2018). Upon their biosynthesis in the cytosol, flavonoids are rapidly accumulated into vacuoles or other cellular compartments through vesicle trafficking, membrane transporters or glutathione S-transferase (GST)-mediated transport (Zhao 2015). In addition to transport of flavonoids into vacuoles mediated by ATP-binding cassette (ABC) transporters (Francisco et al. 2013), there is also H^+^-dependent transport mechanism in grapevine (Gomez et al. 2011; Kuang et al. 2019). Vacuolar transport of anthocyanins and other metabolites, such as malate, directly or indirectly depends on the transmembrane pH gradient generated by both tonoplast H^+^-ATPases and H^+^-pyrophosphatases (V-PPase) (Shiratake and Martinoia 2007). Three V-ATPase subunits (VIT_03s0038g00790, VIT_18s0001g01020, VIT_04s0008g02460) and two V-PPases (VIT_11s0118g00350, VIT_09s0002g07880) showed strong decrease in protein abundance in HT berries, and this may impair the efficient transport of anthocyanins into the vacuole. Finally, the translational repression of several GST proteins (VIT_01s0026g01340, VIT_01s0011g01900, VIT_08s0040g03100, VIT_19s0093g00310, VIT_07s0104g01800) observed in heat-stressed berries during ripening could have an impact on fruit secondary metabolism as well (Perez-Diaz et al. 2016).

#### Harmful effects of heat stress on carbohydrate and energy berry metabolism

A drastic negative effect of HT was observed for proteins assigned to primary carbohydrate metabolism (**Supplementary Table S9**). Almost all the DAPs related to the following processes were under-accumulated after HT: major CHO metabolism (6 out of 8 DAPs), glycolysis (8 out of 9 DAPs), fermentation (4 out of 5 DAPs), gluconeogenesis (3 out of 4 DAPs), oxidative pentose phosphate pathway (6 DAPs) and TCA cycle (13 out of 14 DAPs). To better understand the consequences of this HT repressive effect, and to expand beyond the previously reported metabolite changes (Lecourieux et al., 2017), we measured an additional 31 metabolites that are involved in sugar or organic acid metabolism, or respiration. To put the measured metabolites into a metabolic context, a schematic representation of plant central carbon metabolism was created to display the metabolite profiles under HT compared to control conditions, for each developmental stage and each stress duration (**Figure 3, Supplementary Table S10**). A significant effect of the developmental stage (two-way ANOVA, *p* < 0.05) was observed for most quantified metabolites (23 out of 31), in agreement with the previously reported metabolite profiles in developing berries from Cabernet Sauvignon fruit-bearing cuttings (Dai et al. 2013). HT significantly influenced the content of 14 metabolites including shikimate, glycerate, glycerol-3-P (G-3-P), several glycolytic intermediates (fructose 6-phosphate (Fru6P), 3-phosphoglycerate (3-PGA) and pyruvate), fumarate, and several sugar phosphates (glucose 6-P (G6P), trehalose 6-P (T6P), sucrose 6’-P (S6P) and mannose 6-P (M6P)). The well-known accelerated decrease of malate under elevated temperature was also observed although the changes were below the level of significance in an ANOVA test. However, when considered over the 14-day time frame of the experiments, the decrease in malate was statistically significant according to a t-test (**Figure 3**). This agrees with the significantly lower malate content detected previously in HT berries at harvest (Lecourieux et al., 2017).

Metabolism undergoes profound reprogramming during fleshy fruit development (Beauvoit et al. 2018). The contents of primary and secondary metabolites have been investigated in developing grape berries (Dai et al. 2013; Degu et al. 2014; Cuadros-Inostroza et al. 2016; Wang et al. 2017). Using Cabernet Sauvignon fruit-bearing cuttings as in the present work, Dai et al. (2013) conducted a deep analysis of metabolites from central carbon metabolism using an LC-MS/MS approach. They reported the timing of important switches in primary carbohydrate metabolism during grape berry development and discussed the relationship between these metabolite changes and transcriptional and translational modifications within the fruit. Similarly, Wang et al. (2017) reported close correlations between the metabolome and the proteome at the interface of primary and secondary metabolism in developing *V. vinifera* Early Campbell fruits. Down-regulation of carbohydrate production and protein synthesis is a conserved response of plants to abiotic stresses (Cramer et al. 2011). This may be a way for plants to save energy, and reflects a shift from plant growth to mechanisms of protection. However, with the exception of organic acids, the effects of high temperature on berry primary metabolism in general are still poorly described (Serrano et al. 2017). It is well known that warm climatic conditions enhance the degradation of malate in grapevine fruits, leading to a decrease in total acidity (Sweetman et al. 2014). Sugar metabolism is desynchronized from organic acid metabolism in ripening berries at high temperatures (Rienth et al. 2016). The micro-environment plays an active role in controlling the dynamics of primary metabolite accumulation that affect fruit composition. For instance, heating strongly alters the berry concentration of various amino acids, including phenylalanine and γ-aminobutyric acid, and decreases malate content (Lecourieux et al. 2017). Integrating microclimatic parameters and metabolomic analysis, Reshef et al. (2017) reported that solar irradiance affected the overall levels and patterns of accumulation of sugars, organic acids, amino acids and phenylpropanoids, across the grape cluster. The relationship between the diurnal microclimatic changes (radiation and temperature) and the berry metabolic dynamics was recently described (Reshef et al. 2019). The present study provides additional evidence about the harmful effects of heat on berry primary metabolism through the quantification of 31 metabolite intermediates related to sugar accumulation, glycolysis, and the TCA cycle, as well as through the identification of numerous proteins related to the corresponding metabolic pathways whose abundance is affected by HS. Among the 31 metabolites quantified, 14 were significantly impacted upon HT, with reduced contents in warmed berries for all of these **(Figure 3, Supplementary Table S10)**. Some intermediates were more affected when the HT was applied during the green stage (glycerate, fumarate), and others during ripening (T6P and S6P), whereas the remaining ones were negatively impacted at all three developmental stages (pyruvate, shikimate, G3P, 3-PGA, G6P, Man6P, F6P). Reduced malate contents were also observed under HS, including the green stage (**Figure 3**). These primary metabolite profiles coincide with a dramatic down-regulation of numerous enzymes related to carbohydrate and energy metabolism in heat-treated berries (48 out of 56 DAPs, **Supplementary Table S9**), and more than 70% of these DAPs were repressed during the ripening period. Using a Cabernet Sauvignon cell suspension culture subjected to HS (42°C), George et al. (2015) observed a similar decline in proteins related to sugar metabolism. Our results highlighted changes in many metabolic pathways in grape berries exposed to heat, possibly due to the inhibitory effect of elevated temperature on metabolic enzyme activities from these pathways.

Several enzymes from the pathways of sugar and sugar-phosphate metabolism were less abundant in heated berries, including a fructokinase (FRK, VIT_14s0006g01410) and two hexokinases (HXK, VIT_18s0001g14230, VIT_11s0016g03070). The decrease in these enzymes may contribute to the lower concentration in sugar phosphates (T6P, S6P, G6P and F6P) under heat, since the hexoses (glucose and fructose) resulting from cleavage of imported sucrose can be converted to hexose phosphates by FRK and HXK in the cytosol (Stein and Granot 2018). In plants, trehalose 6-phosphate (T6P) acts as a signal of sucrose availability connecting plant growth and development to its metabolic status, and it was reported that T6P levels can exhibit dynamic responses to environmental cues (Figueroa and Lunn 2016). An increase in T6P content was observed in berries from veraison on, but this increase was lower under heat (**Figure 3, Supplementary Table S10**). In some plants, T6P inhibits the SnRK1 Ser/Thr protein kinase, in turn contributing to a reduction in anthocyanin content (Baena-Gonzalez et al. 2007). It is thus tempting to speculate that the increase in T6P concentration from veraison onwards contributes to anthocyanin accumulation through repression of SnRK1, and that this regulatory process is less effective under heat. However, the roles of T6P as a signalling molecule and the SnRK1/T6P pathway during the berry ripening process and in HT response remain to be demonstrated. In sink organs such as developing berries, sucrose cleavage is carried out either by invertase (INV) to yield glucose and fructose, or by sucrose synthase (SUS) to yield UDPG and fructose. Our proteomic analysis revealed a strong accumulation of a SUS isoform after HT at veraison (SUS2, VIT_05s0077g01930). However, glucose, fructose and sucrose contents were not differentially altered by heat treatment (**Figure 3, Supplementary Table S10**), whereas a delay in the increase in total soluble solids (TSS) content was observed in developing berries after heat exposure during the green stage (Lecourieux et al. 2017). Accumulation of SUS may be a sign of hypoxia resulting from increased respiration rate under high temperature conditions, thus consuming oxygen faster than it can diffuse into the bulky grape berry tissue. Such as *AtSUS1* and *AtSUS4* in Arabidopsis, *SUS* genes are described as hypoxic responsive genes and a role for sucrose synthase as part of the acclimation mechanism to anoxia in dicots was proposed (Santaniello et al., 2014).

The present study also revealed the lower abundance of many glycolytic enzymes (9 DAPs) in warmed berries, mostly during ripening, together with the reduced content of some glycolytic intermediates (Fru6P, 3-PGA, pyruvate). A similar decrease in abundance of glycolytic enzymes or glycolytic intermediates was also reported for some grape varieties including Cabernet Sauvignon, when berries ripen under normal growth conditions (Giribaldi et al. 2007; Dai et al. 2013; Martinez-Esteso et al. 2013). Thus, the consequence of heat stress on the glycolytic pathway remains unclear since the glycolytic flux might have been maintained of even increased at high temperature although the decrease of some glycolytic-related DAPs. Disturbing the glycolytic pathway may impair the generation of energy (ATP), reducing equivalents (NADH) and various intermediates required for the production of amino acid, lipids and secondary metabolites. For example, it has been reported that cell energy, reducing power and α-ketoglutarate availability are potential drivers for flavonoid biosynthesis in grape cells (Soubeyrand et al. 2018).

The TCA cycle is a central metabolic hub since it generates energy, reducing power and carbon skeletons. Several enzymes from the TCA cycle were repressed in heated berries but consequences on the corresponding TCA intermediates were not obvious within the time frame of the analysis. The transcript levels of MDH, which catalyzes a reversible reaction between oxaloacetate and malate, and ME which oxidizes malate to pyruvate, were reported to increase during grape berry ripening. This higher expression might contribute to the decline of malate concentration after veraison (Martinez-Esteso et al., 2011; Wang et al, 2017). In our study, however, heat exposure reduced protein amount of both enzymes in ripening fruits (MDH: VIT_10s0003g01000, VIT_19s0014g01640, VIT_00s0373g00040; ME: VIT_15s0046g03670, VIT_11s0016g03210), therefore questioning their precise role in hastening the malate decrease under heat. The amount of a phosphoenolpyruvate carboxylase (PEPC, VIT_19s0014g01390) was also reduced by HT at the green stage. This may contribute to lower the amount of malate under heat before veraison (**Figure 3**), since the first step of anaplerotic malate synthesis involves the carboxylation of PEP by PEPC (Sweetman et al. 2009). However, the loss of malate during ripening has been attributed to increased degradation rather than decreased synthesis pre-veraison (Ruffner et al. 1976; Sweetman et al. 2014). Accordingly, PEPC activity decreases in pre-veraison berries with day heating, and malate content did not correlate with changes in PEPC activity (Sweetman et al. 2014). The PEPC activity is highly regulated by reversible protein phosphorylation and mono-ubiquitination, which affects its sensitivity to feedback inhibition by malate (Ruiz-Ballesta et al. 2016). Thus, changes in the phosphorylation and/or the ubiquitination status of PEPC might be more important than changes in total PEPC protein abundance. Energy-consuming processes such as gluconeogenesis may also be impacted in ripening berries exposed to HT, with three down-regulated enzymes (MDH, PEPCK and isocitrate lyase). Lower MDH amounts would reduce the accumulation of oxaloacetate (OAA), required for PEP formation through phosphoenolpyruvate carboxykinase activity (PEPCK). In normal growth conditions, both PEPCK activity and transcript abundance are increased in berries after veraison (Walker et al. 2015). In ripening grapes, PEPCK may function in gluconeogenesis when malate released from the vacuole exceeds the demand from other malate-consuming processes (Walker et al. 2015).

#### Heat stress promotes changes in lipid metabolism and cell wall remodelling

The functional category “Lipid metabolism” was particularly affected in heat-exposed berries since all the lipid-related proteins (20 DAPs) identified in our work were negatively regulated after HT (**Supplementary Table S2**). Moreover, we also observed that warmed berries contain lower amount of glycerol 3-phosphate, precursor of the membrane glycerolipids (**Figure 3**). These lipid-related DAPs are involved in fatty acid (FA) synthesis, FA elongation or FA degradation, among which were the following enzymes: acetyl-CoA carboxylase (ACC, VIT_18s0001g04980, VIT_11s0065g00360), glycerol-3-phosphate acyltransferase (GPAT, VIT_15s0046g02400), beta-ketoacyl-CoA synthase (KCS, VIT_13s0067g03890) and phosphoinositide phosphatase (SAC, VIT_04s0044g00030). ACC proteins catalyse the carboxylation of acetyl-CoA to form malonyl-CoA during fatty acid synthesis (Sasaki and Nagano 2004). The cytosolic ACCase contributes to the synthesis of very-long chain fatty acids, with a role in membrane stability and biosynthesis of flavonoids. Its lower abundance in warmed berries may negatively impact the flavonoid accumulation upon veraison. The heat-repressive effects observed at the protein level may also lead to a deep remodelling of the berry protection system. Particularly, the down-regulation of some proteins such as GPAT and KCS may disturb the berry protective barrier, namely the cuticle, since these two proteins belong to families involved in cutin and wax biosynthesis, respectively (Guo et al. 2016; Petit et al. 2016). The strongest translational repression was observed for a protein from the SAC (suppressor of actin) family (log_2_FC: −6.22) when the HS was applied during the green stage. SAC proteins are phosphatases that hydrolyze polyphosphoinositides, and roles in stress responses, cell wall formation, vacuolar trafficking and vacuolar morphology and function have been demonstrated in plants (Novakova et al. 2014). Among the known effects of high temperatures, one of the most relevant is the alteration of membrane-linked processes due to modifications in membrane fluidity and permeability (Niu and Xiang 2018). Heat stress decreases the total content of lipids and glycolipids (Escandon et al. 2018). To preserve membrane integrity under elevated temperatures, plants can regulate the ratio of bilayer to non-bilayer-forming lipids (Welti et al. 2007). Plant thermotolerance might also be improved by reducing the membrane content of unsaturated fatty acids or by accumulating more saturated fatty acids prior to HS, thus helping to maintain optimal membrane fluidity (Higashi et al. 2015; Escandon et al. 2018). Accordingly, we found that two desaturase proteins (VIT_05s0094g00750, VIT_00s0499g00030) were down-regulated in warmed berries. Membrane components may also be involved in initiating key signalling pathways that lead to the appropriate plant responses to heat (Niu and Xiang 2018). Consequences of the heat-induced lipid metabolism changes for berry development and quality remain to be determined.

The cell wall forms a dynamic boundary for plant cells and plays important roles during grape berry development, governing the potential for cell expansion during fruit growth and modulating the texture of mature berries through the depolymerisation and the solubilisation of cell wall polymers (Goulao et al. 2012b). Cell wall remodelling also constitutes an important component of plant responses to HS that maintain overall function and growth, although the cell wall factors contributing to the acquisition of plant thermotolerance remain largely unknown (Wu et al. 2018). Our previous transcriptomic analysis highlighted the strong effect of HT on genes involved in cell wall homeostasis, and especially on enzymes contributing to cell wall expansion and loosening such as xyloglucan endo-transglycosylase and expansins (Lecourieux et al. 2017). At the protein level, most of the 25 DAPs related to the category “cell wall” displayed lower amounts in heated berries, except for a few proteins among which were two pectin methylesterases (PME, VIT_07s0005g00730, VIT_11s0016g00290) and two polygalacturonases (PG, VIT_02s0025g01330, VIT_08s0007g08330) that accumulate in ripening berries exposed to heat. Both PME and PG are implicated in pectin disassembly and fruit softening process, but it is difficult to assess the impact of changes in these four proteins, since both enzymes belong to large families, with 36 PMEs and 60 PGs being annotated in the reference grape genome (Goulao et al. 2012a). However, there is increasing evidence that cell wall-modifying enzymes are involved in plant responses to HS, and PMEs appear to be key players in the process (Wu et al. 2018). Similar observations were made in fruits of other species exposed to high temperature, such as strawberry (Langer et al. 2018).

### Identification of proteins with a potential role in berry responses to heat

Global warming represents a substantial risk for the sustainability of wine production in many regions. Plant breeding combined with molecular plant biotechnology has the potential to deliver stable yields and quality under warmer climate conditions, but this requires the identification of key traits of tolerance to heat (Torregrosa et al. 2017). Our proteomic study revealed 54 DAPs related to the functional category “Stress” and mainly belonging to the “abiotic/heat stress” cluster (**Supplementary Table S7**; 35 out of the 54 “stress” DAPs). Most of the proteins associated with the “heat stress” category were over-accumulated, mainly at veraison stage, and all were members of the heat shock protein (HSP) family. Looking for potential protective mechanisms activated in berries exposed to HT, we paid particular attention to up-regulated proteins previously described in the HS responses and thermotolerance in other plant species (**Table 2; Supplementary Tables S7 and S12**). We also listed some proteins that increased in abundance under HT and thus have a putative role in stress tolerance. These proteins (**Table 2**) are potential markers for developing or selecting grape varieties that are better adapted to warmer climate or the extreme heat waves that are expected due to climate change.

**Table 2.**
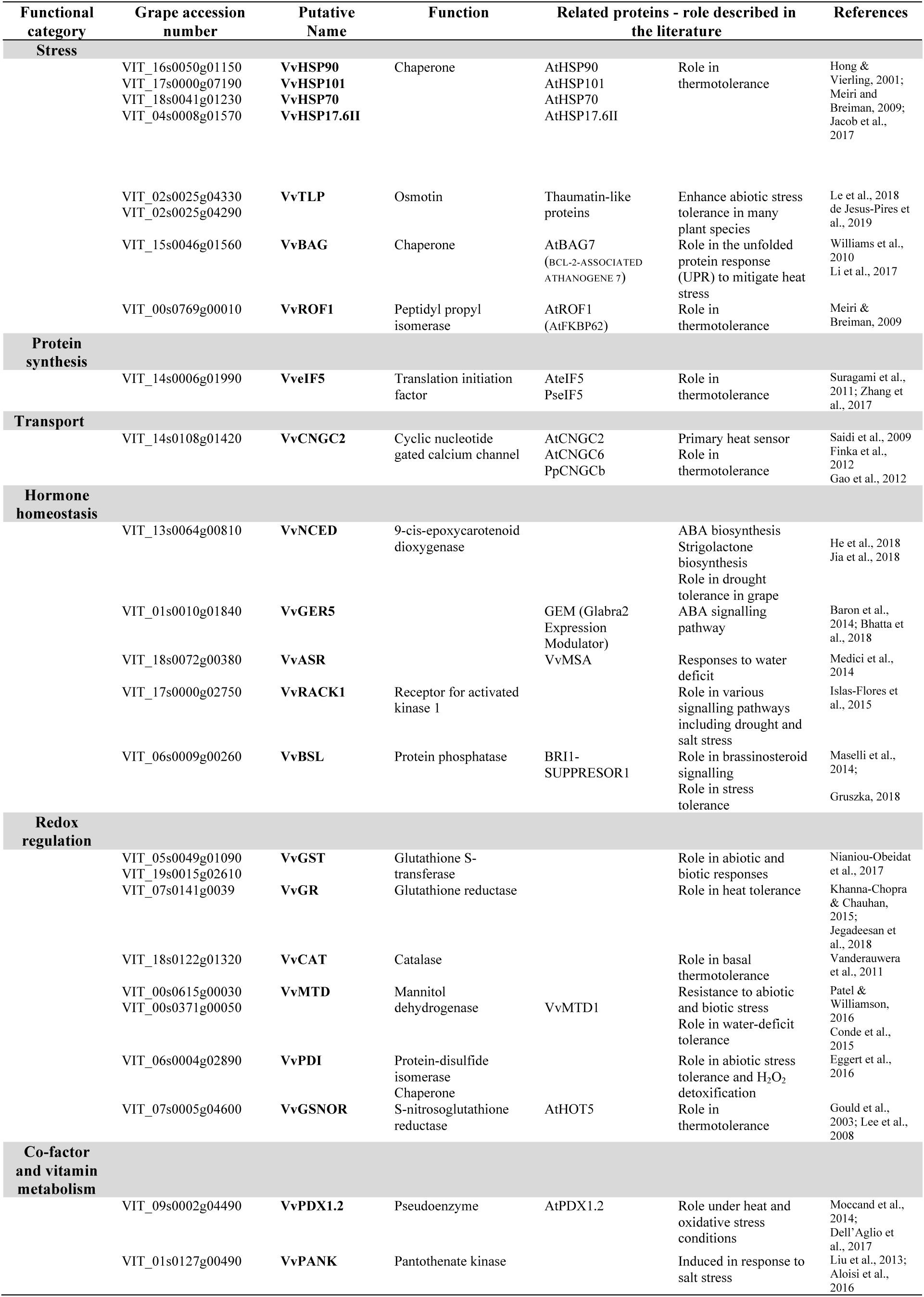
Deregulated proteins with potential role in heat-induced berry responses

Cabernet Sauvignon berry can improve its HS response abilities by elevating the levels of particular proteins associated with the maintenance of protein homeostasis (synthesis, folding and degradation). HSP/chaperones are considered as powerful buffers against environmental stress including HS. Their chaperone functions include not only protein folding, but also signalling and protein targeting and degradation (Bokszczanin and Fragkostefanakis 2013). Data collected over the last decades suggest that members of the HSP/chaperone network are valuable targets for engineering multiple stress resistance in crops (Jacob et al. 2017). Among the 30 up-regulated HSPs in heat-exposed berries, four of these were increased irrespective of the fruit developmental stage (HSP70, HSP90, HSP101, HSP17.6II), and all of these were described as required for thermotolerance in various plant species (**Table 2**). Another up-regulated protein under HS corresponds to a Bcl-2-associated athanogene isoform (BAG). Plant BAGs are multifunctional proteins that regulate cell development and contribute to cytoprotection from pathogen attack and abiotic stress (Doukhanina et al. 2006). In particular, AtBAG7 was described to be a key player of the heat-induced unfolded protein response (UPR) pathway that alleviates the detrimental effects of accumulation of misfolded proteins (Li et al. 2017). VvROF1 abundance was also increased in HS conditions. In Arabidopsis, AtROF1 encodes a peptidyl prolyl isomerase catalysing the *cis*/*trans* isomerization of peptide bonds N-terminal to proline residues within polypeptide chains. AtROF1 is transcriptionally induced by HT and is involved in acquired thermotolerance through the interaction with HSP90.1 and HSFA2 (Meiri and Breiman 2009). We also observed that two thaumatin-like proteins (VvTLPs) from the osmotin sub-family strongly accumulated in heat stressed berries. The grape genome contains 33 VvTLPs (Yan et al. 2017), and this family is known to be involved in biotic and abiotic stress responses including grapevine (Liu et al. 2010). Osmotins have been shown to have multiple functions in enhancing stress tolerance in many plant species, although the mechanisms by which osmotins mediate plant response to abiotic stress are not well established (de Jesus-Pires et al. 2019). Finally, a strong increase in the translation initiation factor VveIF5 was detected in heated berries (**Table 2**). The role of eIF5 in thermotolerance was described in Arabidopsis and pea (Suragani et al., 2011; Zhang et al., 2017).

How plants sense and transduce the HT signal is still an important topic to be addressed. It was reported that phytochromes function as plant thermosensors (Jung et al. 2016). Transduction of the HT signal involves various players (Li et al. 2018), including members of the plasma membrane cyclic nucleotide gated calcium channel family (CNGC). In Arabidopsis, CNGC2 and CNGC6 were described as heat sensors, contributing to the Ca^2+^ signalling induced by HT and leading to the onset of plant acquired thermotolerance (Finka et al. 2012). Whereas *CNGC6* gene is ubiquitously expressed in Arabidopsis (Gao et al. 2012), our results revealed that the protein abundance of the grape ortholog VvCNGC2 increased after berry exposure to heat, underlining the need for *de novo* synthesis of this protein to achieve its role in thermotolerance.

Integration of elevated temperature, signal transduction and the appropriate stress response is mediated, at least partially, by plant hormones (Ku et al. 2018). Several proteins related to hormone homeostasis were seen to be more abundant in heat-exposed berries (**Table 2**). Four abscisic acid (ABA)-related DAPs were up-regulated in the present study. The first DAP is a 9-cis-epoxycarotenoid dioxygenase (NCED). This enzyme catalyses the first committed step in ABA biosynthesis (Tan et al. 1997) and NCED overexpression improves drought tolerance in grape (He et al. 2018). At the transcript level, a mixed picture has emerged for the *NCED* isogenes in HS grape berries (Carbonell-Bejerano et al. 2013; Rienth et al. 2014a; Lecourieux et al. 2017). The second ABA-related DAP was GER5/GRE5, a protein closely related to GEM (GLABRA2 Expression Modulator) and belonging to a subfamily of the GRAM domain-containing proteins (Mauri et al. 2016). In heat-stressed berries, GER5 was up-regulated at both mRNA and protein levels (**Supplementary Table S11**). GER5 and closely related GRAM domain genes (GER1, GEM) were proposed to be required in the reproductive development of Arabidopsis whereas a putative role in abiotic stress responses remains unclear (Bhatta et al. 2018). The third ABA-related DAP is an ABA-, stress-, and ripening-(ASR) induced protein known to play prominent roles in the protection of plants against abiotic stress (Gonzalez and Iusem 2014). For instance, the grape ASR VvMSA was reported to be involved in responses to water deficit (Medici et al. 2014). The last ABA-related DAP is RACK1 (receptor for activated C kinase 1). In rice, this protein positively regulates seed germination by controlling endogenous levels of ABA and H_2_O_2_ (Zhang et al. 2014). More broadly, plant RACK1 proteins regulate various signalling pathways ranging from developmental processes such as seed germination, flowering and leaf initiation, to immune and stress responses against pathogens and environmental stimuli (Islas-Flores et al. 2015). Another phytohormone-related DAP was a brassinosteroid-related protein phosphatase which accumulated in heated berries. In Arabidopsis, this protein is one of a four-member gene family (BSL, for BRI1-SUPPRESSOR1 Like) shown to participate in brassinosteroid signalling (Maselli et al. 2014), and potentially involved in stress tolerance (Gruszka 2018).

Heat stress is accompanied by oxidative stress, which may trigger adverse effects through oxidative damage of cell membranes and important macromolecules (Bokszczanin and Fragkostefanakis 2013). Therefore, redox regulation and signalling have been recognised as crucial mechanisms for acute responses to HT (Katano et al. 2018). Our proteomic analysis of HT berries highlighted several up-regulated DAPs with antioxidant properties, including two glutathione S-transferases (GSTs), a glutathione reductase (GR, VIT_07s0141g0039), one catalase, a protein-disulfide isomerase (PDI), an aldo-keto reductase (AKR), two mannitol dehydrogenases (MTD) and an *S*-nitrosoglutathione reductase (GSNOR). Glutathione S-transferases belong to a well-characterized detoxification enzyme family, catalyzing the conjugation of the reduced glutathione tripeptide (GSH) to electrophilic substrates (Kumar and Trivedi 2018). GSTs quench reactive molecules through the addition of GSH and protect the cell from oxidative damage. Accordingly, plant GSTs play an important role in abiotic and biotic stress responses (Nianiou-Obeidat et al. 2017). The two up-regulated GSTs were observed at veraison but an opposite pattern was seen at ripening with five GST isoforms being strongly repressed at the protein level. This GST down-regulation observed under HT may negatively impact the accumulation efficiency of flavonoids into the vacuolar compartment of ripening berries. Indeed, GSTs function as non-enzymatic carriers in intracellular transport, allowing the accumulation of flavonoid-GSH conjugates into vacuoles (Petrussa et al. 2013). The GST-mediated transport of anthocyanins and proanthocyanidins has been established in grape as well (Perez-Diaz et al. 2016). Among the four other heat-affected proteins, GR catalyses the reduction of glutathione disulfide (GSSG) to GSH, and its role in heat tolerance was described in different plant species (Jegadeesan et al. 2018). The ROS detoxifying enzyme catalase is considered as an acclimation protein required for basal thermotolerance (Vanderauwera et al. 2011), whereas *PDI* encodes an oxidoreductase enzyme belonging to the thioredoxin superfamily. PDI can be considered as a folding enzyme since it catalyses the formation of correct disulfide bridges between the cysteine residues in proteins to maintain ER-homeostasis (Park and Seo 2015). In potato, StPDI1 was found to interact with a sucrose transporter (StSUT1) (Krugel et al. 2012), and a role in abiotic stress tolerance and H_2_O_2_ detoxification was suggested (Eggert et al. 2016). Recently, the heterologous over-expression of a *Methanothermobacter thermautotrophicus* PDI with a chaperone function and disulfide isomerase activity could confer thermotolerance to transgenic rice (Wang et al. 2018). The aldo-keto reductase (AKR) superfamily contains proteins able to reduce a broad spectrum of substrates, ranging from simple sugars to potentially toxic aldehydes. Plant AKRs are involved in diverse metabolic reactions including reactive aldehyde detoxification, biosynthesis of osmolytes, secondary metabolism and membrane transport. These proteins may also confer multiple stress tolerance (Sengupta et al. 2015), including heat stress tolerance as reported in rice (Turoczy et al. 2011). Two mannitol dehydrogenases (MTD) increased in abundance in grape berries exposed to HT. The role of mannitol and by extension, its catabolic enzyme MTD, is well established in resistance to both biotic and abiotic stresses. Several studies suggest that mannitol protects plants from various abiotic stresses by acting as an osmoprotectant and by quenching the damaging ROS (Patel and Williamson 2016). In grapevine, VvMTD1 was described as involved in water-deficit stress tolerance (Conde et al. 2015). Finally, GSNOR has an important function in the homeostasis of NO (nitric oxide), which is considered as a broad-spectrum anti-stress molecule when acting at low concentration in plants. Whereas NO was described to accumulate in tobacco exposed to HT (Gould et al. 2003), the key role of GSNOR1-mediated de-nitrosylation in thermotolerance was revealed by the characterization of Arabidopsis *hot5* (sensitive to hot temperatures) knock-out mutants (Lee et al. 2008). Taken together, these up-regulated and redox-related proteins may play a role in signalling as well as in preventing harmful effects of ROS and reactive nitrogen species in berries under HS conditions.

Finally, our proteomic work highlighted the heat-induced accumulation of two proteins related to vitamin metabolism, namely VvPDX1.2 and VvPANK (**Table 2**). The pseudoenzyme PDX1.2 (Pyridoxine biosynthesis 1) is a noncatalytic homolog of the PDX1 subunit of the vitamin B_6_ biosynthesis protein machinery. Transcriptionally regulated by HsfA1, PDX1.2 serves to stabilize the catalytic PDX1s under HT, thereby maintaining vitamin B_6_ homeostasis in plants (Dell’Aglio et al. 2017). AtPDX1.2 was reported as essential for plant growth under heat and oxidative stress conditions (Moccand et al. 2014). The second up-regulated protein corresponds to a pantothenate kinase (PANK). Pantothenate (known as vitamin B5) is the key precursor of CoA and acyl carrier protein, which are essential co-factors for many metabolic enzymes (Raman and Rathinasabapathi 2004). The PANK converts pantothenate to phosphopantothenate, using ATP as phosphate donor. PANK activity plays a critical role in regulating intracellular coenzyme A (CoA) levels in bacteria and animals but little is known about the role of this enzyme in plants (Ottenhof et al. 2004). It was suggested that an increased level of CoA, through an increased PANK activity, may be responsible for improved plant growth and stress resistance (Rubio et al. 2008).

### Conclusion

Global warming represents a substantial risk for the sustainability of wine production in many regions. Plant breeding combined with molecular plant biotechnology have the potential to deliver stable yields and quality under warmer climate conditions, but this requires the identification of key traits of tolerance to heat. To this end, we combined omics approaches to decipher the molecular components and processes affected in developing Cabernet Sauvignon berries exposed to heat. In the present study, 592 differentially abundant proteins were identified in heated berries among which some may play a substantial role in heat tolerance. Our data also show that there is not a straightforward correlation between the heat-modulated berry transcriptome and proteome, indicating the major involvement of post-transcriptional regulation under heat stress. These pronounced changes contribute on the one hand to generate fruit damaging effects, including disruption of carbohydrate and secondary metabolism, and on the other hand to mobilize key heat-signalling players as well as protective molecules such as antioxidants and chaperones (**Figure 5**). In summary, these findings help elucidate heat-triggering effects on developing grape berries and identify genes and proteins that potentially control heat acclimation in grapevine. The functional characterization of some putative candidates is in progress.

**Figure 5.**
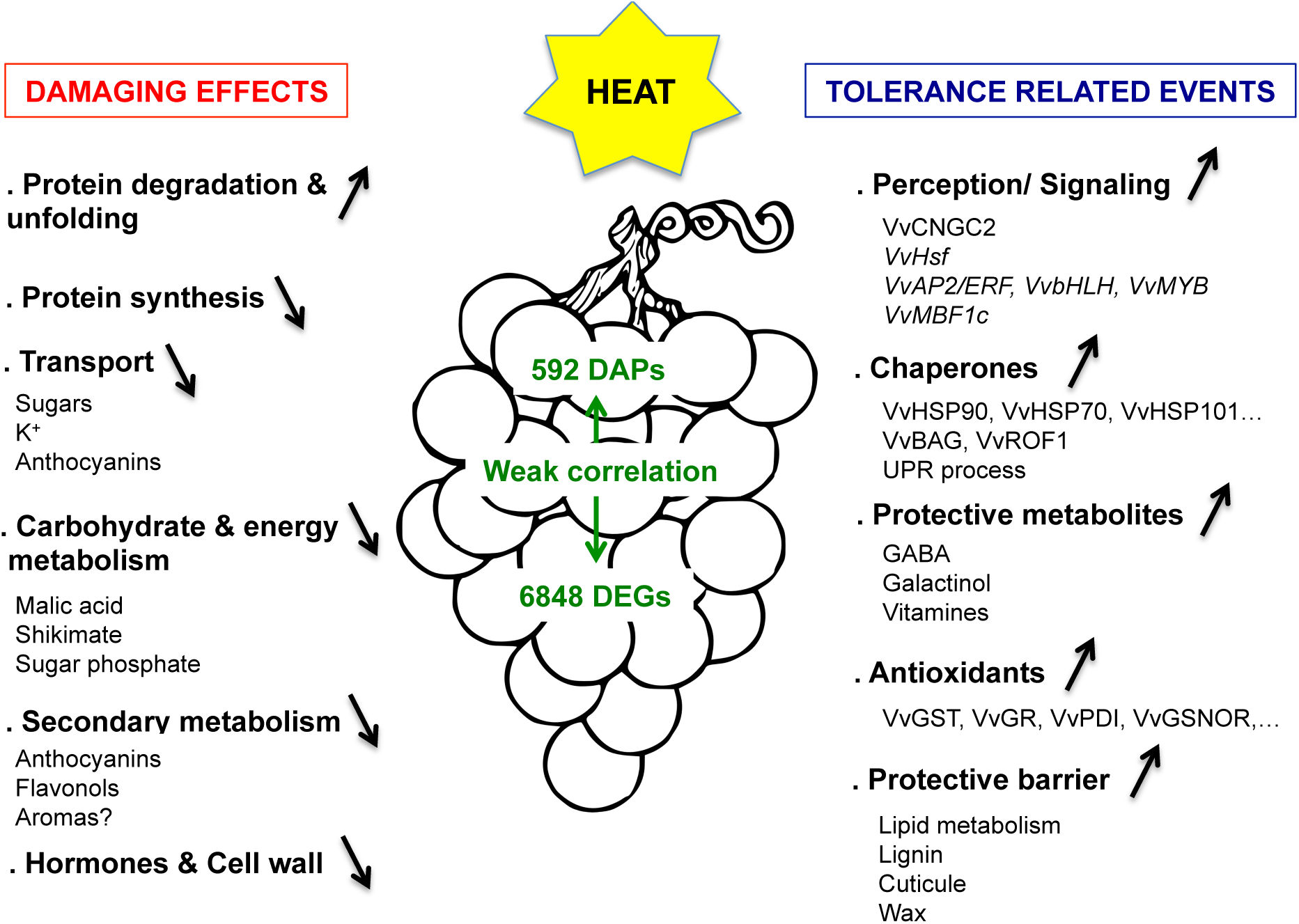
Schematic overview of heat stress effects on developing Cabernet Sauvignon berries. This illustration summarizes the main findings deduced from transcriptomic, proteomic and metabolomics analyses (this work and (Lecourieux et al. 2017)). Up and down arrows reflect the accumulation levels of the corresponding transcripts, proteins or metabolites in heated berries.

## MATERIAL AND METHODS

### Plant material, temperature and sampling

Elevated temperature (+ 8°C) was directed on clusters of *V. vinifera* Cabernet Sauvignon fruiting cuttings. The HT was applied at three different berry developmental stages and maintained throughout the light period (7:00 am to 7:00 pm) until harvesting. To separate short- and long-term responses, berries were collected after 1 and 7 days of treatment. Details of the experimental protocol and sampling were provided in Lecourieux et al. (2017). The same pools of deseeded berries were used for both the transcriptomic analysis reported in Lecourieux et al. (2017) and the proteomic analysis reported here.

### Tissue preparation and total protein extraction

Each biological replicate contained 25 pooled berries underwent independent protein extraction using a protocol adapted from Lücker et al. (2009) and Martinez-Estezo et al. (2011). Briefly, deseeded berries were ground to a fine powder in a mortar with liquid nitrogen and 3 g of frozen ground tissue were used to prepare a total protein extract. Pectins and pigments were removed after incubation of the frozen tissues in 25 mL of a cold (−20°C) ethyl acetate: ethanol (1 : 2 (v/v) for 30 min at −20°C with repeated vortexing. The protein extract was precipitated by centrifugation for 3 min at 21000 × *g* at 4°C, and a second ethyl acetate: ethanol wash was performed in a similar way. The pellet was next extracted twice with cold (−20°C) 100% acetone by vortexing and centrifuging, as before. The pellet was transferred to a mortar and dried in a fume hood until complete acetone evaporation. The dried pellet was finely ground after addition of 1/3 vol of sand, and washed with 1.5 mL chilled 10% (w/v) trichloroacetic acid (TCA) in acetone. This washing step was repeated seven times. This was followed by two washes with aqueous 10% (v/v) TCA, two with chilled 80% (v/v) acetone, and drying at 4 °C. Each washing step included incubation at −20°C for 20 min followed by centrifugation for 3 min at 21000 × *g* at 4°C. Subsequently the dried pellet was resuspended in 500 µL protein extraction buffer containing: 0.7 M sucrose, 0.1 M KCl, 0.5 M Tris-HCl (pH 7.5), 50 mM EDTA, 1% PVPP, 1% DTT and protease inhibitor cocktail (Roche, Germany), and incubated at room temperature (RT) for 30 min with intermittent vortexing. Then, an equal volume of Tris-saturated phenol (pH 7.5) was added, the sample was incubated at RT for 30 min with vortexing every 5 min. After centrifugation at 15,000 × *g* at RT for 40 min, the phenol phase in the upper layer was transferred into a new 2-mL Eppendorf tubes and the remaining aqueous lower phase was re-extracted with another 500 µL Tris-saturated phenol (pH 7.5). The two phenol phases were combined and washed twice with an equal volume of phenol washing buffer (0.7 M sucrose, 0.1 M KCl, 0.5 M Tris (pH 7.0, 50 mM EDTA, 1% DTT, and protease inhibitor cocktail). Proteins contained in the organic phase were precipitated overnight at 4°C with 5 volumes of 0.1 M ammonium acetate in methanol. Precipitated proteins were washed three times in 0.1 M ammonium acetate in methanol and twice in ice-cold 80% (v/v) acetone. All these precipitating and washing steps included incubation for 30 min followed by centrifugation at 15,000 × *g* at 4°C for 40 min. Finally, proteins were dissolved in 200 µL freshly prepared solubilization solution (7 M urea, 70 mM SDS, 20 mM DTT) and quantified according to the Bradford method (Bradford, 1976) with bovine serum albumin as standard.

### Sample preparation and protein digestion

Thirty µg of each protein sample (three biological replicates for each condition; 12 conditions in total: three stages x two treatments x two treatment durations) were solubilized in Laemmli buffer and were separated by SDS-PAGE in a 10% (w/v) acrylamide gel. After colloidal blue staining, each lane was horizontally divided into four parts of equal length and each part was subsequently cut in 1 mm x 1 mm gel pieces. Gel pieces were destained in 25 mM ammonium bicarbonate dissolved in 50% (v/v) acetonitrile (ACN), rinsed twice in ultrapure water and dehydrated in 100% ACN for 10 min. After ACN removal, gel pieces were dried at room temperature, covered with proteolysis solution (containing 10 ng/µl trypsin dissolved in 40 mM NH_4_HCO_3_ and 10% (v/v) ACN), rehydrated at 4 °C for 10 min, and finally incubated overnight at 37 °C. The gel pieces were then incubated for 15 min in 40 mM NH_4_HCO_3_ and 10% (v/v) ACN at room temperature with rotary shaking. The supernatant was collected, and an H_2_O/ACN/formic acid (47.5:47.5:5 by volume) extraction solution was added and gel slices incubated for 15 min. The extraction step was repeated twice. Supernatants were pooled and concentrated in a vacuum centrifuge to a final volume of 40 µL. Digests were finally acidified by addition of 2.4 µL of 5% (v/v) formic acid and the resulting peptide mixtures stored at −20 °C until analysis.

### nLC-MS/MS analysis

Peptide mixtures were analyzed on an Ultimate 3000 nanoLC system (Dionex, Amsterdam, The Netherlands) coupled to a nanospray LTQ-Orbitrap XL mass spectrometer (ThermoFinnigan, San Jose, USA). Ten microliters of peptide mixtures were loaded onto a 300-µm-inner diameter x 5-mm C_18_ PepMap^TM^ trap column (LC Packings) at a flow rate of 30 µL/min. The peptides were eluted from the trap column onto an analytical 75-mm id x 15-cm C18 Pep-Map column (LC Packings) with a 5– 40% linear gradient of solvent B in 105 min (solvent A was 0.1% formic acid in 5% ACN, and solvent B was 0.1% formic acid in 80% ACN). The separation flow rate was set at 200 nL/min. The mass spectrometer operated in positive ion mode at a 1.8-kV needle voltage and a 39-V capillary voltage. Data were acquired in a data-dependent mode alternating a Fourier transform mass spectrometry (FTMS) scan survey over the range m/z 300–1700 and six ion trap MS/MS scans with CID (Collision Induced Dissociation) as activation mode. MS/MS spectra were acquired using a 3-m/z unit ion isolation window and a normalized collision energy of 35%. Only +2 and +3 charge-state ions were selected for fragmentation. Dynamic exclusion duration was set to 30s.

### Database search and MS data processing

Data were searched using the SEQUEST algorithm (supported by the Proteome Discoverer 1.4 software; Thermo Fisher Scientific Inc.) against a subset of the 2012.01 version of UniProt database restricted to *Vitis vinifera* Reference Proteome Set (29,817 entries). Spectra from peptides higher than 5000 Da or lower than 350 Da were rejected. The search parameters were as follows: mass accuracy of the monoisotopic peptide precursor and peptide fragments was set to 10 ppm and 0.8 Da respectively. Only *b*- and *g*-ions were considered for mass calculation. Oxidation of methionines (+16 Da) was considered as a variable modification. Two missed trypsin cleavages were allowed. Peptide validation was performed using the Percolator algorithm (Käll et al. 2007), and only “high confidence” peptides were retained corresponding to a 1% false positive rate at the peptide level.

### Label-Free Quantitative Data Analysis

Raw LC-MS/MS data were imported into the Progenesis LC-MS 4.0 (Non Linear Dynamics) software. Data processing includes the following steps: (i) Features detection, (ii) Features alignment across the six samples for each part, (iii) Volume integration for 2-6 charge-state ions, (iv) Normalization on total protein abundance, (v) Import of sequence information, (vi) ANOVA test at the peptide level and filtering for features p <0.05, (vii) Calculation of protein abundance (sum of the volume of corresponding peptides), (viii) ANOVA test at protein level and filtering for features p <0.05. Only non-conflicting features and unique peptides were considered for calculation at the protein level. Quantitative data were considered for proteins quantified by a minimum of two peptides. The mass spectrometry proteomics data have been deposited to the ProteomeXchange Consortium via the PRIDE (Perez-Riverol et al. 2019) partner repository with the dataset identifier PXD014693.

Protein abundance changes between control and heat-exposed fruits were considered to be significant when a minimum fold change of 1.5 (log_2_FC > 0.58 or < −0.58) was reached, with a significance threshold of *p* < 0.05. Only proteins identified in all three biological replicates were considered for quantification. Significantly affected protein categories based on the MapMan Ontology (Usadel et al., 2005) were identified using a Chi-square test. MapMan mappings for the Cribi 12X grapevine genome were based on closest homologs regarding to the Arabidopsis thaliana genome. Protein annotations were taken from Grimplet et al. (2012).

### Primary metabolism quantification

Deseeded berries were ground to a fine powder in liquid nitrogen and aliquots (15–20 mg) of frozen powder were extracted with chloroform/methanol as described by Lunn et al. (2006). Primary metabolites were measured by LC-MS/MS as described by Lunn et al. (2006), with modifications as described in Figueroa et al. (2016) and Dai et al. (2013).

## ACKNOWLEDGMENTS

The authors would like to thank Christelle Renaud and Ghislaine Hilbert for metabolite quantifications. For the production of the fruiting cuttings and technical assistance during greenhouse experiments, we express our gratitude to Jean Pierre Petit, Nicolas Hoquard and Guillaume Pacreau.

## AUTHOR CONTRIBUTIONS

DL, FL and PP designed the research; DL oversaw the research; PP and DL performed the greenhouse experiments; FL, JC, and DL carried out the protein extraction; CK performed the bioinformatics analysis; RF, DL and JEL did the metabolic analysis; SC and MB did the metabolic analysis, FL, CK, and DL analysed and interpreted the data; DL and FL drafted the manuscript; SD, JEL, LW and EG critically revised the manuscript. All authors read and approved the final manuscript.

## SUPPORTING INFORMATION

**Figure S1:** MapMan GO categories impacted at the transcriptomic and proteomic levels in heated grape berries

**Table S1:** All identified proteins with calculated differentials and corresponding statistics

**Table S2:** List and information about the differentially abundant proteins between control and heat stress conditions

**Table S3:** Differentially abundant proteins between control and heat stress condition at green stage

**Table S4:** Differentially abundant proteins between control and heat stress condition at veraison stage

**Table S5:** Differentially abundant proteins between control and heat stress condition at ripening stage

**Table S6:** List of “Protein metabolism” related DAPs

**Table S7:** List of “Stress” related DAPs

**Table S8:** List of “Secondary metabolism” related DAPs

**Table S9:** List of “carbohydrate and energy metabolism” related DAPs

**Table S10:** Effects of the HT and developmental stages on individual metabolites,

**Table S11:** List of overlapping DEGs and DAPs in Cabernet Sauvignon berries exposed or not to HT

**Table S12:** DAPs with potential role in thermotolerance.

