## Supplemental Figure S1 for "Proteomic and metabolomic profiling underlines the stage- and time-dependent effects of high temperature on grape berry metabolism"

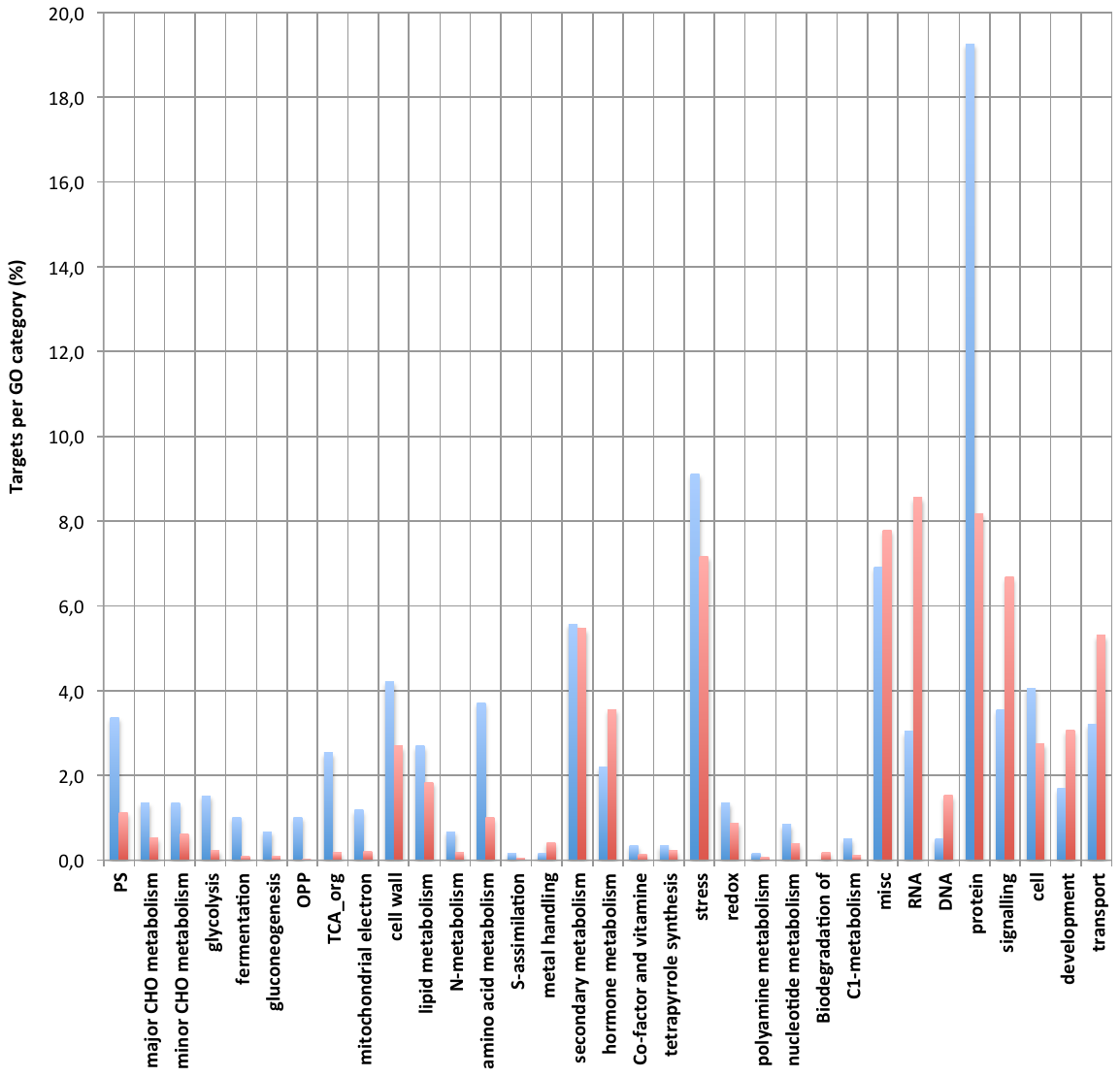

**Supplementary Figure 1: MapMan GO categories impacted at the transcriptomic (red bars) and proteomic (blue bars) levels in heated grape berries.** Percentages of differentially abundant genes or proteins (targets) across the 6 experimental conditions are provided on the y-axis for each category.
